# Individual movement variability magnitudes are predicted by cortical neural variability

**DOI:** 10.1101/097824

**Authors:** Shlomi Haar, Opher Donchin, Ilan Dinstein

**Author notes:** **Corresponding author:** Shlomi Haar. **Author Contributions:** S.H., O.D., and I.D., Conception and design, Interpretation of data, Drafting and revising the article; S.H., Data acquisition and analysis. **Conflict of Interest:** The authors declare no competing financial interests.

## Abstract

Humans exhibit considerable motor variability even across trivial reaching movements. This variability can be separated into specific kinematic components such as extent and direction, which are thought to be governed by distinct neural processes. Here, we report that individual subjects exhibit different magnitudes of kinematic variability, which are consistent (within individual) across movements to different targets and regardless of which arm (right or left) was used to perform the movements. Simultaneous fMRI recordings revealed that the same subjects also exhibited different magnitudes of fMRI variability across movements in a variety of motor system areas. These fMRI variability magnitudes were also consistent across movements to different targets when performed with either arm. Cortical fMRI variability in the posterior-parietal cortex of individual subjects predicted their movement-extent variability. This relationship was apparent only in posterior-parietal cortex and not in other motor system areas, thereby suggesting that individuals with more variable movement preparation exhibit larger kinematic variability. We, therefore, propose that neural and kinematic variability are reliable and interrelated individual characteristics that may predispose individual subjects to exhibit distinct motor capabilities.

## INTRODUCTION

Intertrial variability is a fundamental characteristic of human movements (e.g., Harbourne and Stergiou, 2009). Variability of specific kinematic components such as movement extent and movement direction is thought to be governed by independent neural processes (van Beers, 2009; Gordon et al., 1994a; Krakauer et al., 2000) according to the demands of the examined motor task (Latash et al., 2007; Todorov, 2004). While kinematic variability is detrimental for movement accuracy, it is thought to be critical for motor learning (e.g., Braun et al., 2009; Herzfeld and Shadmehr, 2014; Teo et al., 2011; Wilson et al., 2008; Wu et al., 2014).

Intertrial variability is also a fundamental characteristic of neural activity, which is apparent in the variable timing and amplitude of neural responses across trials containing an identical stimulus or task (e.g., Churchland and Abbott, 2012; Dinstein et al., 2015; Faisal et al., 2008; Sauerbrei et al., 2015; Stein et al., 2005). As with kinematic variability, intertrial neural variability also seems to be important for motor learning as demonstrated in studies with songbirds (Kao et al., 2005; Olveczky et al., 2011; Woolley and Kao, 2015) and primates (Mandelblat-Cerf et al., 2009). Given that neural activity generates behavior, one may expect that intertrial variability in the activity of specific neural populations would generate corresponding intertrial variability in specific kinematic components of movement (e.g., movement extent and/or direction).

Studies that have examined the potential relationship between neural and kinematic variability have proposed three alternative theories. The first theory proposed that kinematic variability during visually guided movements is mostly explained by variability in sensory neural populations. For example, intertrial variability in the initial speed of smooth-pursuit eye movements can be explained by variability in the estimation of target speed in MT neurons (Osborne et al., 2005; for review, see Lisberger and Medina, 2015). In contrast, the second theory has proposed that kinematic variability during reaching movements is generated by variable preparatory (motor planning) activity of premotor and primary motor neurons (Churchland et al., 2006). Finally, the third theory has suggested that kinematic variability is caused by neural and neuro-muscular variability during actual movement execution (van Beers, 2009; van Beers et al., 2004). Taken together, these studies suggest that distinct neural variability sources are correlated with kinematic variability under different experimental conditions, which include the sensory-motor requirements of the examined motor task (e.g., smooth-pursuit ocular movements versus reaching movements) and the temporal structure of the task (e.g., imposing a delay between movement planning and execution). Notably, most of the electrophysiology studies described above have focused on the relationship between movement velocity variability (rather than movement extent or direction) and neural variability.

In the current study we examined several outstanding questions regarding kinematic variability, neural variability, and their potential relationship in humans: 1. Do individual subjects exhibit consistent magnitudes of kinematic variability regardless of the movements that they are performing (e.g., when using right or left arm)? 2. Do individual subjects exhibit consistent magnitudes of neural variability regardless of the movements that they are performing? 3. If so, are between-subject differences in kinematic variability explained by differences in neural variability in specific sensory and/or motor brain areas? Answering these questions is critical for establishing that individual subjects exhibit characteristic kinematic and neural variability magnitudes that may predispose them to exhibit particular motor learning capabilities while also adding new insights regarding the potential relationship between neural variability and kinematic variability.

To answer the questions above and relate the findings with the existing behavioral and electrophysiology literature we quantified intertrial variability of movement direction, peak velocity, and extent across slice (out-and-back) reaching movements. These movements were performed to four peripheral targets with either right or left arm on a touch screen while brain activity was recorded with fMRI. We then quantified fMRI response variability in the primary motor, premotor, parietal, and visual brain areas of each subject and examined whether it was possible to predict between-subject differences in kinematic variability according to neural variability magnitudes in specific brain areas. Note that in our study all movements were performed without visual feedback to preclude the potential influence of neural variability associated with visual input.

## RESULTS

### Intertrial kinematic Variability

Subjects exhibited considerable intertrial kinematic variability in their slice (out-and-back) movements to each of the four targets (Figure 1B). We focused our analyses on three kinematic components: direction (at end-point) and extent, which are commonly reported in behavioral studies (van Beers, 2009; Gordon et al., 1994a; Krakauer et al., 2000), and peak velocity, which is commonly reported in electrophysiology studies (Churchland et al., 2006; Cisek, 2006). Note that movement extent and peak velocity are mutually dependent, because peak velocity scales with increasing target distance (Gordon et al., 1994b).

**Figure 1.**
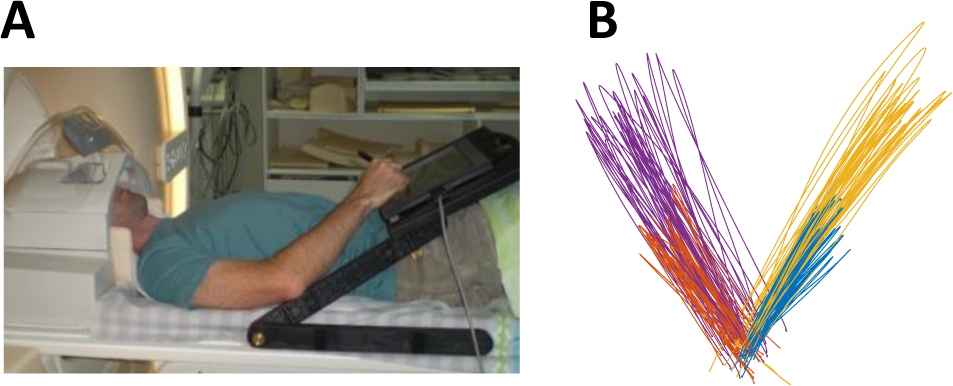
**(A)** *Experimental setup.* **(B)** Representative example of movement paths of one subject. Different direction (at end-point) and extent, which colors represent slice movements to the four targets.

In line with previous findings, we found that the variance of movement extent and peak velocity grew with the mean (correlation across subjects: r = 0.35 and r = 0.53 respectively, averaged across targets and arms). To examine differences in intertrial variability not explained by differences in the mean, we used the coefficient of variation (CV). In contrast, mean movement direction was not correlated with its standard deviation across trials (r < 0.1). There was, therefore, no reason to normalize this measure, so we used the standard deviation (SD) across trials to quantify movement direction variability.

When examining each of the kinematic components separately, individual subjects exhibited consistent magnitudes of intertrial variability across movements to different targets (Figure 2A&B). Thus, subjects who were, for example, more variable in their movement extents to one target tended to be more variable in their movement extents to all other targets. We quantified this by computing the mean Pearson correlation coefficients across all target pairs for movements performed with the right arm (r = 0.29, 0.41, and 0.39 for movement direction, extent, and peak velocity respectively, q(FDR) < 0.001) and left arm (r = 0.46, 0.58, and 0.40 for movement direction, extent, and peak velocity respectively, q(FDR) < 0.001). Significant correlations were also evident when comparing the variability magnitudes of each kinematic component across arms (Figure 2C). For example, subjects with more variable movement extents in right arm movements exhibited more variable movement extents in left arm movements as well (r = 0.63, 0.68, and 0.54 for movement direction, extent, and peak velocity, p < 0.001).

**Figure 2.**
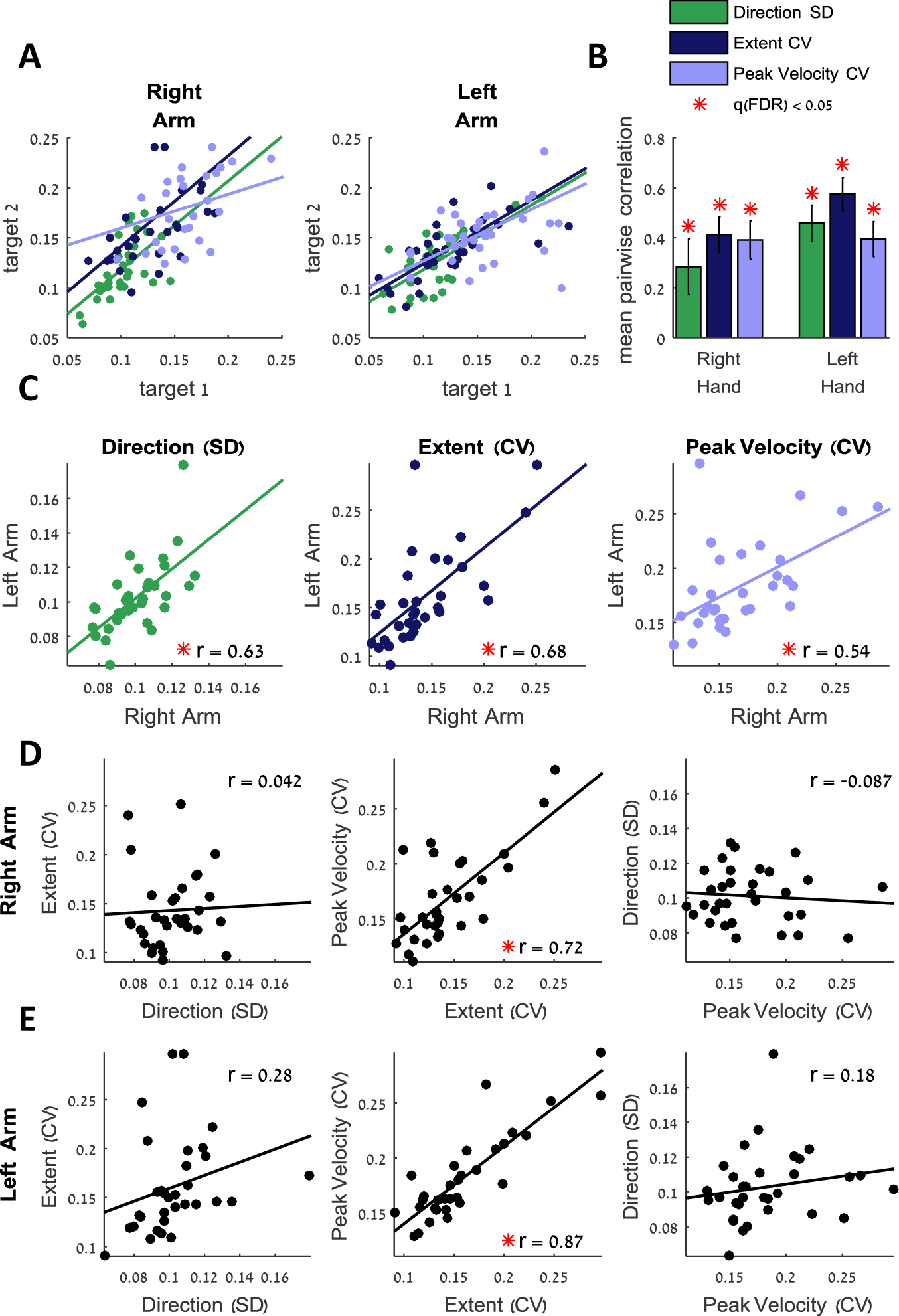
Kinematic variability correlations. (**A**) Movement direction (standard deviation, SD), movement extent (coefficient of variation, CV), and peak movement velocity (CV) of left and right arm movements toward a pair of targets. Each data point represents the variability of movements of one subject to the two targets. (**B**) Means and SEM of the Pearson correlations of the variability across all pairs of targets. Significant correlations are marked with asterisks. (**C**) Scatter plots of the kinematic variability, averaged across targets, of the right and left arms. Each data point represents variability of movements of a single subject. (**D,E**) Scatter plots of the kinematic variability, averaged across targets, of the right (**D**) and the left (**E**) arms. For all scatter plots: data points represent different subjects; lines represent linear fits. Significant correlations are marked with red asterisks.

In line with previous reports (Gordon et al., 1994b), intertrial variability of movement extent and peak velocity were strongly correlated in movements of the right arm (r = 0.72, p < 0.001; Figure 2D) and left arm (r = 0.87, p < 0.001; Figure 2E), but variability of movement extent and movement direction (right arm: r = 0.04, p = 0.41; left arm: r = 0.28, p = 0.07) or peak velocity and movement direction (right arm: r = −0.09, p = 0.69; left arm: r = 0.18, p = 0.16) were not. Thus, individuals who exhibited large movement extent and peak velocity variabilities did not necessarily exhibit large movement direction variability and vice versa.

### Intertrial fMRI variability

All subjects exhibited robust fMRI responses during the execution of movements, which enabled us to identify five cortical ROIs that are commonly examined in motor system studies (Figure 3): Primary Motor Cortex (M1), dorsal premotor cortex (PMd), ventral premotor cortex (PMv), supplementary motor area (SMA), superior parietal lobule (SPL), and inferior parietal lobule (IPL). In addition to the motor ROIs we also identified ROIs in early visual cortex (Vis), dorsolateral prefrontal cortex (dlPFC), and outside the brain (OOB).

**Figure 3.**
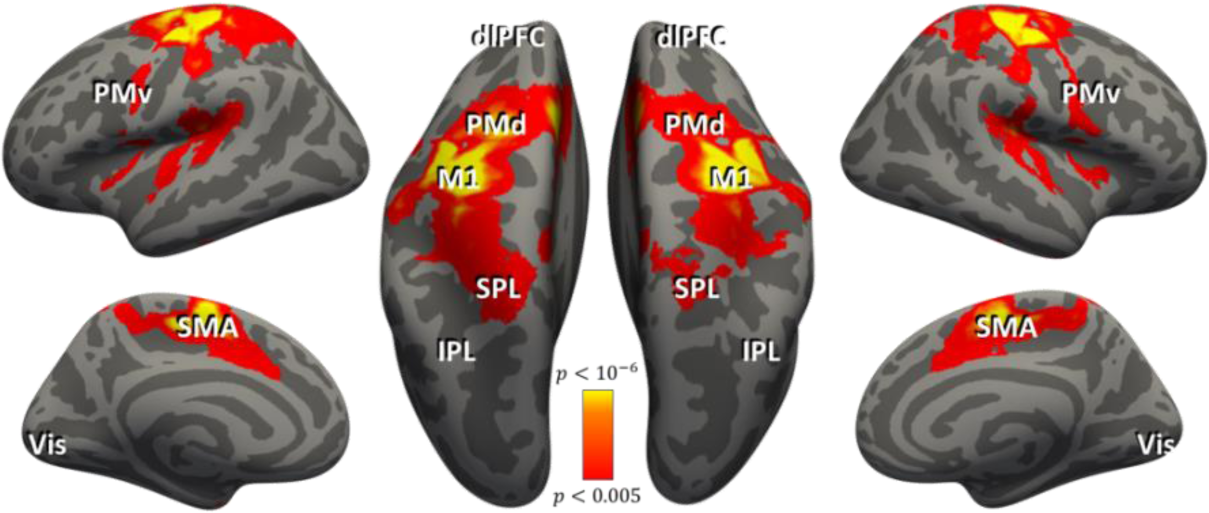
Regions of interest. Cortical areas that exhibited larger responses during arm movement are shown in red/orange. Results calculated across all subjects (random-effects GLM) and displayed on inflated hemispheres of a template brain. The general locations of the selected ROIs are noted (actual ROIs were anatomically and functionally defined in each subject – see Methods). ROIs: Primary motor cortex (M1), dorsal premotor cortex (PMd), ventral premotor cortex (PMv), supplementary motor area (SMA), inferior parietal lobule (IPL), superior parietal lobule (SPL), dorsolateral prefrontal cortex (dlPFC), and the visual cortex (Vis).

We then quantified intertrial fMRI variability in each of the ROIs, separately for each subject, in the following manner: First, we estimated the hemodynamic response function (HRF) in each ROI for each target by averaging the fMRI responses across all movements to that target (Figure 4A). We then used the target-specific HRF in a GLM analysis to estimate a response amplitude/beta-value for each trial/movement in the experiment (Figure 4B). Note that using a target-specific HRF enabled us to compute single trial responses/beta-values relative to the mean HRF of each subject. This approach discounted potential between-subject differences in the mean amplitude and shape of individual HRFs. Finally, we quantified intertrial fMRI variability by computing the standard deviation across beta-values for each of the targets (Figure 4B&C).

**Figure 4.**
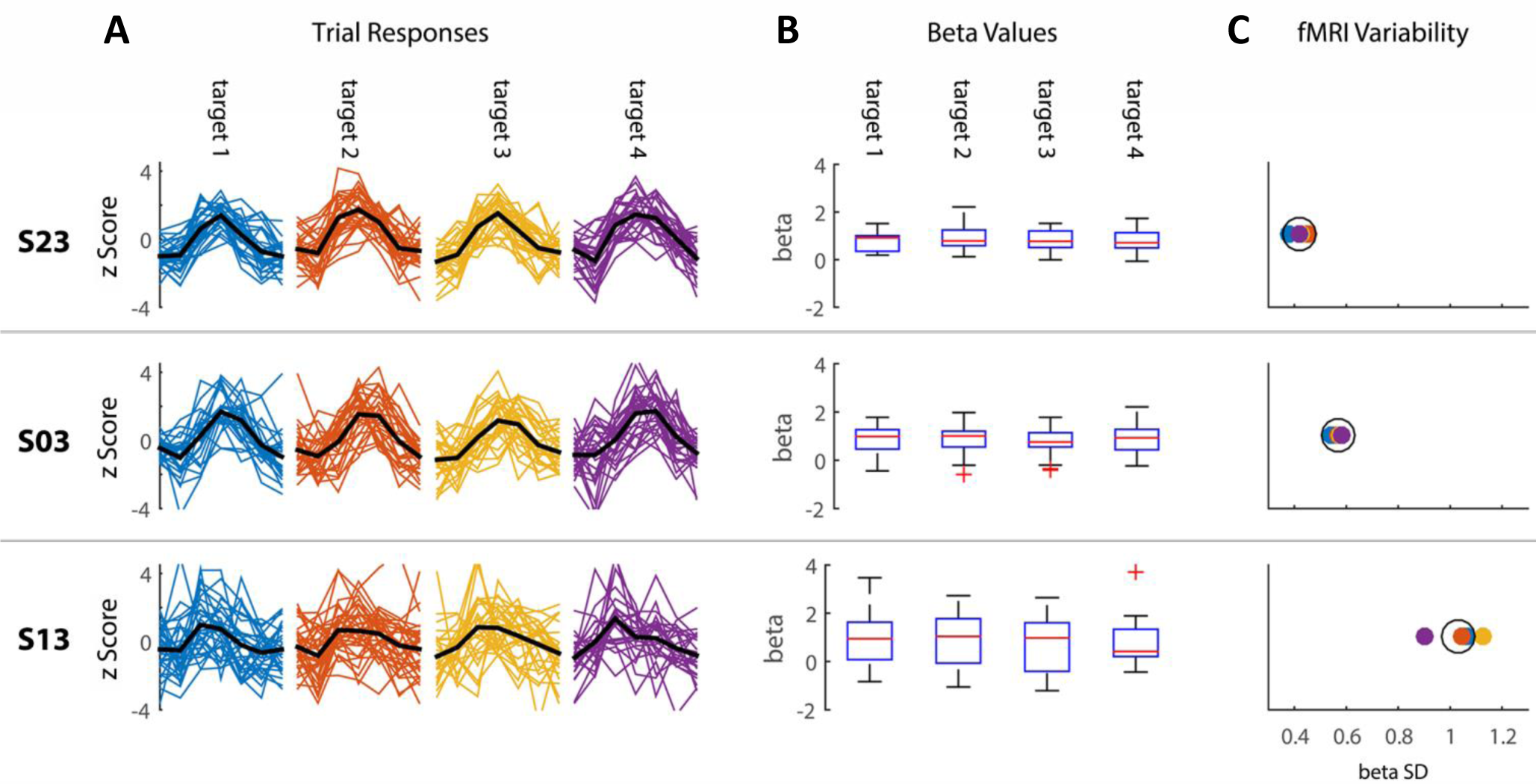
fMRI Variability. Examples of intertrial fMRI variability as quantified in left Ml of 3 subjects during right arm movements. (**A**) Single trial fMRI responses/segments from left M1 are presented in z-scored units; color coded for the different targets, mean HRF across trials (used in the GLM analysis) is presented in black. (**B**) Boxplots demonstrating the distributions of beta-values per target. (**C**) Standard deviation (SD) across beta values for each target (color code is the same as in A). The mean SD across targets is represented by the black circle. Each row represents data from a single subject.

Intertrial fMRI variability was correlated across all pairs of targets in most of the examined ROIs (Figure 5A). Hence, subjects who exhibited more variable brain responses when moving to one target also exhibited more variable brain responses when moving to other targets. During right arm movements all ROIs in the left hemisphere except dlPFC, and all ROIs in the right hemisphere except PMd and dlPFC, exhibited significant pair-wise correlations across targets (r > 0.32, q(FDR) < 0.05). Correlations in the dlPFC and out of brain (OOB) ROIs were not significant (r < 0.26, q(FDR) > 0.1). Taken together, these findings demonstrate that correlation in fMRI variability magnitudes across targets was specific to cortical ROIs that were activated by the task. Note that early visual cortex was activated by the presentation of a visual cue that instructed subjects about the target location at the beginning of each trial (5 seconds before the go cue). Similar results were also apparent for left arm movements (data not shown).

**Figure 5.**
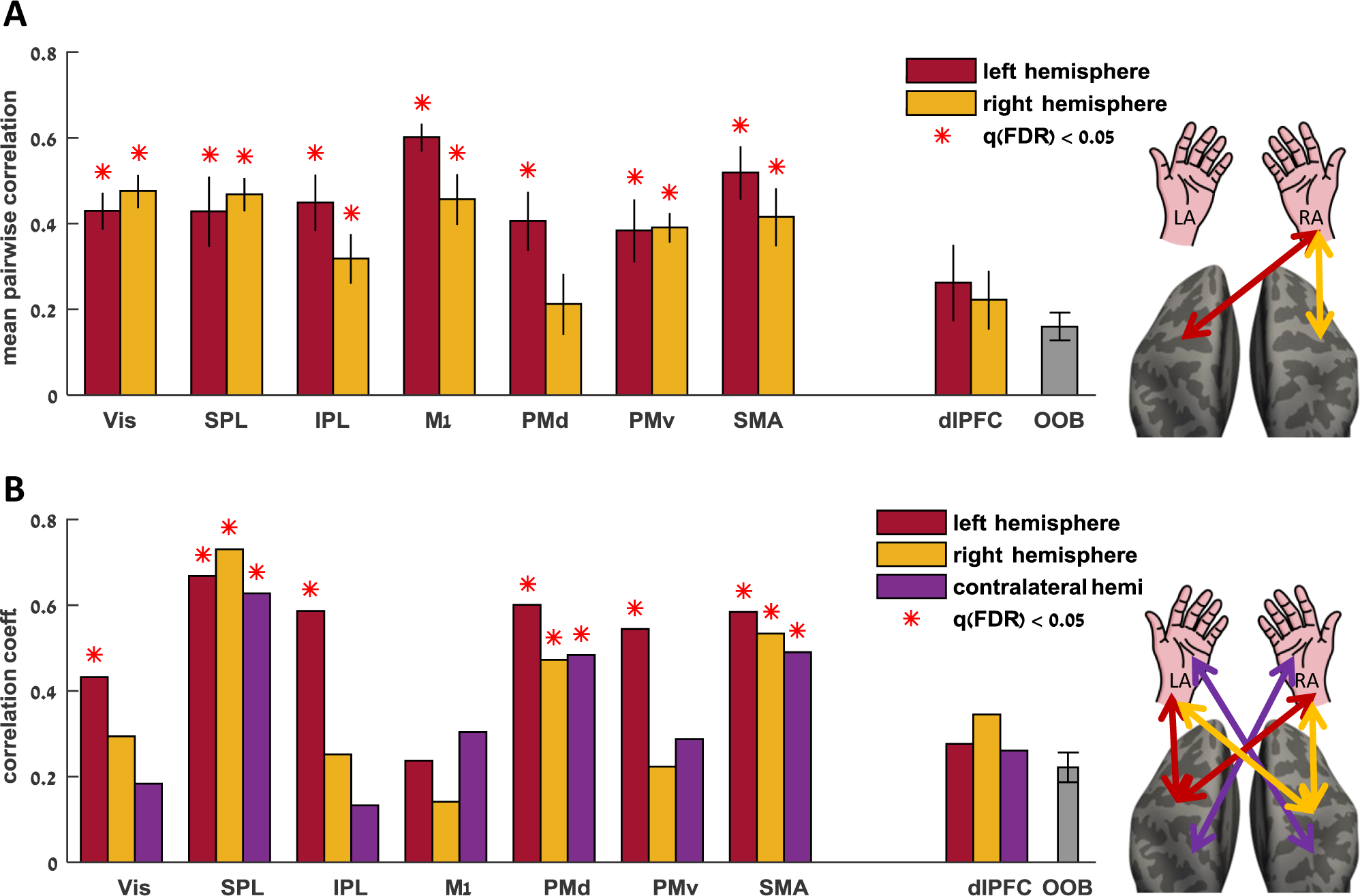
Cortical variability correlations. (**A**) fMRI variability magnitudes during right arm movements were correlated across all target pairs. Mean pair-wise correlation coefficients are presented for each left hemisphere (i.e., contralateral, red) and right hemisphere (i.e., ipsilateral, yellow) ROI. (**B**) fMRI variability magnitudes were correlated across right and left arm movements in left hemisphere ROIs (red), right hemisphere ROIs (yellow) and in contra-lateral ROIs (purple). Significant correlations are marked with red asterisks.

Individual magnitudes of fMRI variability were also significantly correlated across right and left arm movements in many of the examined motor ROIs (Figure 5B). This was evident in all ROIs in the left hemisphere (r > 0.43, q (FDR) < 0.05; Figure 5B, red bars) except for M1 and dlPFC, and in the SPL, PMd, and SMA in the right hemisphere (r > 0.47, q(FDR) < 0.05; Figure 5B, yellow bars). In addition, fMRI variability magnitudes were significantly correlated across left and right arm movements in contralateral SPL, PMd, and SMA ROIs (r > 0.48, q(FDR) < 0.05; Figure 5B, purple bars). This means that, for example, variability in left PMd during right arm movements was significantly correlated with variability in right PMd during left arm movements. Note that consistent fMRI variability across targets and hands was mostly apparent in parietal and prefrontal motor areas, yet was entirely absent in M1. Correlations in the dlPFC and out of brain (OOB) ROIs were not significant (r < 0.33, q(FDR) > 0.09). This demonstrates that consistent fMRI variability differences across subjects were not due to differences in scanner measurement noise across subjects. Such scanner noise differences would be apparent in multiple ROIs and even in ROIs located outside the brain.

### Relationship between kinematic and fMRI variability

Subjects with larger intertrial fMRI variability in the IPL exhibited larger intertrial extent variability (Figure 6). We examined to what extent between-subject differences in kinematic variability could be explained by fMRI variability measures across bilateral ROIs using partial least squares regression. Intertrial fMRI variability in the IPL explained 24% (q(FDR) = 0.004) of the between-subject differences in extent variability, 15% of the variability in the peak velocity, and 8% of the variability in movement direction. The IPL was the only ROI where there was a significant relationship between fMRI variability magnitudes and any of the kinematic variability measures. In contrast, intertrial fMRI variability in M1 explained only 2%, 5%, and 4% (q(FDR) > 0.5) of the between-subject differences in direction, extent, and peak velocity variability respectively. Correlations were not significant in all the control ROIs (dlPFC and out of brain, R^2^ < 8%, q(FDR) > 0.2).

**Figure 6.**
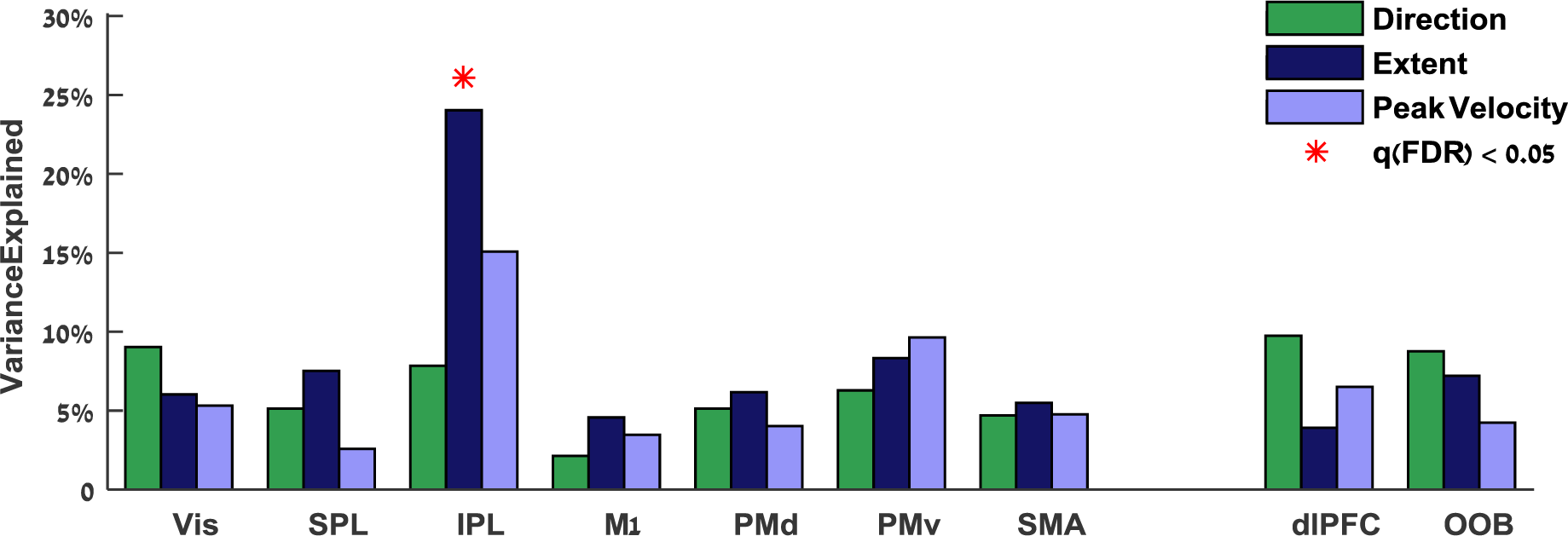
Kinematic Variability explained by fMRI Variability. Multiple regression was performed between fMRI variability magnitudes in each pair of ROIs (right and left hemispheres) and variability magnitudes of three kinematic variables: direction (green), extent (dark blue), and peak velocity (light blue). Significant explained variance is marked with red asterisks (q(FDR) < 0.05).

### Searchlight analysis

To examine the spatial selectivity of the cortical-kinematic relationship we performed an additional analysis using a whole-brain searchlight approach (Kriegeskorte et al., 2006). We mapped the correlations between kinematic variability magnitudes and fMRI variability magnitudes across the entire cortical surface, so as not to restrict the analysis to a-priori ROIs. We used a volumetric searchlight cube of 125 functional voxels in the cortical gray matter segmented within the native space of each subject. For each searchlight cube, we calculated the intertrial fMRI variability (as described above for the ROIs) and then registered the resulting variability maps of all subjects to a common inflated brain. We calculated Pearson correlation coefficients to estimate the relationship between intertrial fMRI variability magnitudes and variability magnitudes of each kinematic variable: movement extent, peak velocity, and direction.

This analysis yielded three searchlight maps that revealed complementary results to those described above. We did not find any cortical areas where fMRI variability magnitudes were significantly correlated with variability magnitudes in movement direction or peak velocity. Significant positive correlations, however, were found in bi-lateral inferior parietal cortex when examining movement extent (Figure 7). Note that the searchlight map is highly symmetric across hemispheres and is relatively similar across movements of the right (Figure 7, Red) and left (Figure 7, Blue) arms.

**Figure 7.**
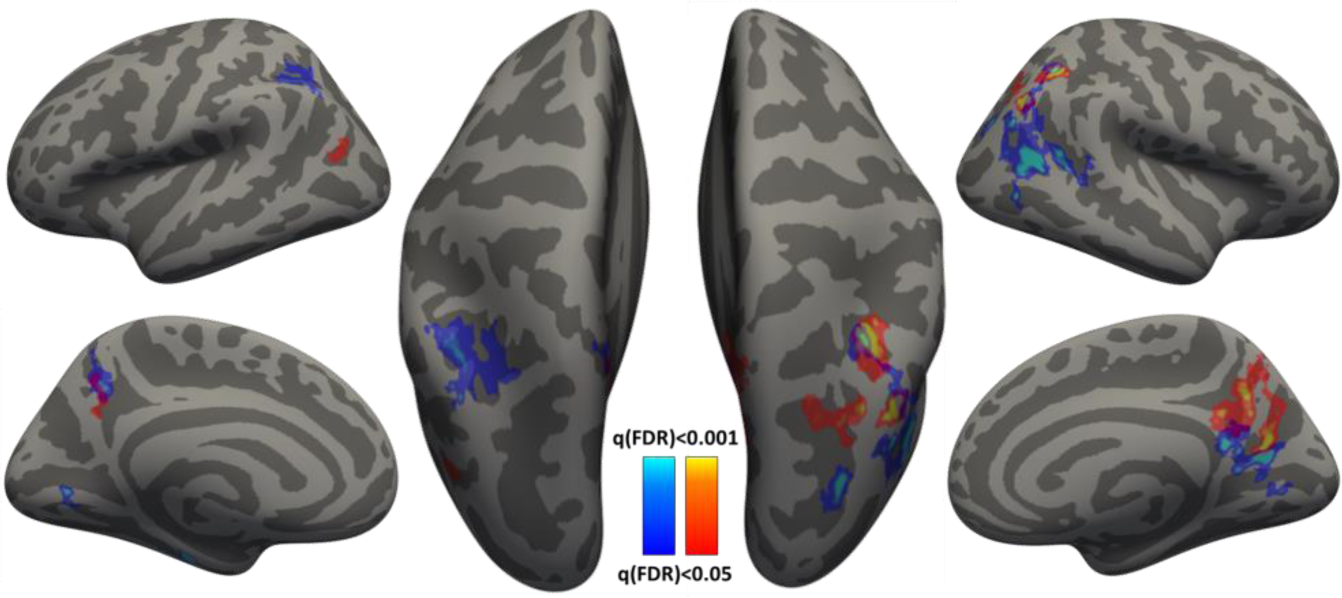
*Searchlight analysis* displaying cortical areas with significant correlations between movement extent variability and fMRI variability across subjects. Results for right (red) and left (blue) arm movements are presented on the inflated cortical anatomy of a single subject. Correlation significance was determined based on a student t-test (FDR corrected).

### Alternative sources of fMRI variability

Between subject differences in fMRI variability can be generated by several non-neural sources that need to be considered. First, previous studies of fMRI variability have reported that the strength of the mean fMRI response was correlated with the magnitude of intertrial variability across subjects (Ferri et al., 2015; He, 2013). To measure intertrial fMRI variability in individual subjects independently of their mean response, we estimated intertrial variability with respect to the mean hemodynamic response function (HRF) apparent in each ROI of each subject (see methods). This enabled us to compute the relative fMRI variability with respect to the actual HRF as opposed to using a canonical HRF that assumes an identical shape and amplitude across subjects. Indeed, when using this method, intertrial fMRI variability was not correlated significantly with mean fMRI response in any of the ROIs (r < 0.15, p > 0.1).

Second, we regressed-out the mean fMRI time-courses of the lateral ventricles and an ROI containing all gray-matter voxels (i.e., “global component”). These time-courses represent fMRI fluctuations that may, in part, be associated with changes in respiration, blood pressure, and other non-neural origins.

Third, head-motion artifacts can generate fMRI variability across trials. To ensure that our results were not generated by head-motion artifacts, we regressed-out estimated head-motion parameters from the fMRI activity of each voxel in the brain before performing the analyses (see methods). Furthermore, we also computed the mean framewise displacement across head-motion parameters (i.e., the mean amount of head motion across samples/TRs) for each subject. We regressed-out individual values of framewise displacement from the fMRI variability magnitudes before examining correlations across targets and/or arms. This ensured that the reported between-subject differences in fMRI variability magnitudes were not generated by underlying differences in head motion across subjects.

## DISCUSSION

Our results reveal that individual subjects exhibit distinct magnitudes of kinematic variability, which are consistent across movements to different locations when performed by either arm. Individual variability magnitudes in movement extent, peak velocity, or direction were strongly correlated across different targets and across arms (Figures 2). This means that an individual who exhibits large movement extent variability to one target is likely to exhibit large movement extent variability to all other targets regardless of the arm that the subject uses to perform the movements.

Analogous findings were also apparent when examining fMRI variability magnitudes of individual subjects (Figures 5). Subjects with larger fMRI variability magnitudes in parietal and pre-motor cortical areas tended to exhibit larger variability regardless of target location or arm used to perform the movements. In contrast, fMRI variability magnitudes in M1 were not consistent across targets or arms. This suggests that cortical variability magnitudes in higher motor system areas are relatively stable over time and across tasks, while cortical variability magnitudes in primary motor cortex represent more transient states that change when subjects perform different movements with different effectors.

The results also revealed a specific relationship between variability magnitudes in one of the kinematic measures, movement extent, and cortical variability magnitudes in one brain area, the IPL. Indeed, fMRI variability magnitudes in the IPL explained 24% of the differences in movement-extent variability across subjects. In contrast, fMRI variability magnitudes in primary motor cortex explained only 5% of between-subject differences in movement-extent variability (Figure 6). The specificity of these results was further validated by a searchlight analysis that revealed significant correlations between the kinematic and cortical variability magnitudes only with respect to movement extent and only in inferior parietal areas (Figure 7). Parietal cortex is thought to play key roles in motor planning, sensory motor mapping, and state estimation (Buneo and Andersen, 2006). We, therefore, suggest that a considerable portion of movement-extent variability is generated by cortical variability associated with movement preparation, rather than cortical variability associated with movement execution.

Note that this is the first study to ever examine the consistency of kinematic variability across targets/hand and relate it with cortical response variability in humans. Contemporary models of motor control and motor learning (Pekny et al., 2015; Wolpert and Flanagan, 2016) emphasize the importance of intertrial variability for motor system flexibility and accuracy. We, therefore, speculate that the stable between-subject differences in cortical and kinematic variability magnitudes described here are likely to predispose individual subjects to exhibit different motor capabilities.

### Neural sources of kinematic variability

Previous theories have suggested that intertrial kinematic variability is predominantly generated by the variable activity of sensory neural populations (Osborne et al., 2005; for review, see Lisberger and Medina, 2015), premotor and primary motor neural populations involved in motor planning (Churchland et al., 2006), or by neuro-muscular variability that characterizes actual movement execution (van Beers, 2009; van Beers et al., 2004). It is entirely possible, however, that different sources of neural variability generate kinematic variability under different experimental conditions, such that behavioral motor variability would embody the sum of multiple neural variability sources (for review see Faisal et al., 2008). With this in mind, neural variability in a particular brain area is likely to explain a certain proportion of kinematic variability. Furthermore, neural variability in different brain areas may generate variability in different kinematic components of movements (e.g., movement extent versus movement direction).

Our results indeed demonstrate that about a quarter of the between-subject differences in movement extent variability are explained by individual neural variability differences in parietal cortex, which is thought to play a dominant role in the planning and preparation of reaching movements (Cohen and Andersen, 2002). While previous electrophysiology studies have reported that variability in M1 and PMd neural activity (during preparation for movement) generates variability in peak movement velocity (Chaisanguanthum et al., 2014; Churchland et al., 2006), our results suggest that stronger relationships between neural and kinematic variability will be evident in parietal brain areas and particularly in IPL (Figure 6&7).

It may seem surprising that correlations between kinematic variability and fMRI variability were remarkably weak in M1 given that M1 is the last brain area in the motor hierarchy that sends out motor commands to the muscles (e.g., Shadmehr and Krakauer, 2008). In humans, however, only 30% to 40% of the axons in the corticospinal tract originate from neurons in the primary motor cortex, while the rest originate from the premotor, supplementary motor, and posterior parietal cortices (Kandel et al., 2013). This means that neural variability in parietal regions may potentially generate kinematic variability downstream of M1, in spinal motor circuits. A potentially interesting analogy can be found in songbirds where a specific nucleus (the lateral magnocellular nucleus of anterior nidopallium – LMAN) has evolved to inject direct neural variability into the motor circuits that control singing – apparently enabling juvenile birds to learn through trial and error (Ölveczky et al., 2011).

Parietal cortex contains neural populations that perform a wide variety of computations that are essential for motor control. Previous studies have focused on aspects of motor planning, sensory motor mapping, and state estimation (Buneo and Andersen, 2006; Churchland et al., 2006; Cohen and Andersen, 2002; Shadmehr and Krakauer, 2008; Wolpert and Ghahramani, 2000). Our analysis localized the peak correlation between the kinematic and the fMRI variability specifically in the IPL of the parietal cortex. Suggested roles of IPL include integration of high-order sensory and motor information in support of high-level motor functions (Fogassi and Luppino, 2005), and conscious motor intentions (Desmurget and Sirigu, 2012). Within all of these frameworks, each with its specific mechanistic focus, variability in the activity of parietal neural populations would generate variability in the kinematics of executed movements. An alternative interpretation of our results, however, might emphasize the sensory roles of parietal cortex. In this case the causality would be reversed such that the measured fMRI variability would be generated by movement variability (and not the other way around). While it is difficult to entirely rule out this option, it is important to note that we did not find significant correlations between fMRI variability magnitudes in primary or secondary somatosensory cortices (Figure 7) and any of the kinematic measures. The selectivity of the results to IPL argues against such a sensory driven explanation of our results.

Finally, it is important to note that we and all previous electrophysiology studies on the topic measured variability only in the kinematics of the movement and not in the dynamics of the executed movement. It is highly possible that intertrial variability in movement dynamics (i.e., muscle activation), which are not necessarily captured in measures of kinematic variability, may be explained by intertrial neural variability in specific brain areas.

### Decomposing neural variability

Neural variability can be decomposed into different spatial and temporal components using measures from different types of neuroimaging and electrophysiological techniques (Dinstein et al., 2015). When studying variability with fMRI, it is possible to simultaneously quantify intertrial variability in multiple different brain areas, but the temporal resolution of this measure is limited by the sluggish nature of the hemodynamic response (Heeger and Ress, 2002). This means that one can estimate a single fMRI response amplitude per trial (in each voxel or ROI) and then quantify intertrial variability across response amplitudes (Figure 4), but estimating intertrial variability in the timing of the response is not possible. Furthermore, since fMRI is not a direct measure of neural activity, but rather a measure of hemodynamic changes over time, intertrial variability in the function of neuro-vascular coupling mechanisms will be an inherent part of the fMRI intertrial variability measure.

This limits the ability to measure neural variability with fMRI and, therefore, limits the ability to relate neural variability and behavioral variability measures. With this in mind it is impressive that we were able to identify a consistent relationship between fMRI variability and movement extent variability which was similarly evident in movements of right and left arm (Figure 7 & 8). We speculate that stronger relationships may be revealed with direct measures of human neural activity such as ECOG recordings in patients who are candidates for epilepsy surgery.

### Hemispheric lateralization

While arm movements are clearly generated and controlled by neural activity in the contralateral hemisphere (Penfield and Boldrey, 1937), we found significant correlations between movement extent variability and neural variability in both the contralateral and ipsilateral hemispheres. We speculate that neural variability in both hemispheres may, therefore, have an impact on the accuracy and reliability of arm movements. Several animal studies have indeed shown that M1 neurons are modulated by ipsilateral arm movements (Cisek et al., 2003; Donchin et al., 1998; Mehring et al., 2003) and could even continuously represent ipsilateral limb position (Ganguly et al., 2009). Furthermore, previous human fMRI studies by us and others found directional tuning of arm movement (Fabbri et al., 2010; Haar et al., 2015) and finger-specific activity patterns across many levels of the cortical motor hierarchy in the ipsilateral hemisphere (Diedrichsen et al., 2013). The ipsilateral neural variability correlations, demonstrated in the current study, emphasize the relationship between the neural activity in the contralateral and the ipsilateral hemisphere, as well as the relevance of the ipsilateral neural activity to motor performance.

### Variability and motor learning

Individual kinematic and neural variability intensities are likely to predispose individual subjects to exhibit different motor learning capabilities. While variability is clearly detrimental for movement accuracy, previous behavioral studies have reported that subjects who exhibited larger task-relevant movement variability were faster learners (e.g., Herzfeld and Shadmehr, 2014; Teo et al., 2011; Wu et al., 2014). Since a considerable portion of the differences in movement variability between subjects can be explained by the differences in their neural variability in IPL, we speculate that individuals with larger neural variability in movement preparation processes may be faster learners. A study relating neural variability with learning rates is, therefore, highly warranted.

### Conclusions

The motor control literature has a long history of examining kinematic intertrial variability across movements to understand how the brain encodes movements and learns to perform new movements. This is the first study to examine how such kinematic variability may be associated with cortical intertrial variability in human subjects. Our results demonstrate that kinematic variability and parietal and pre-frontal cortical variability are stable individual traits, which appear consistently across movements to different targets when performed by either arm. Furthermore, these variabilities are related such that subjects with larger neural variability in IPL exhibited larger movement-extent variability. We believe that these results represent an important first step for understanding how neural variability may generate movement variability in humans and, thereby, predispose individuals to exhibit distinct motor capabilities.

## METHODS

### Subjects

32 right-handed volunteers with normal or corrected-to-normal visual acuity (15 women and 17 men, aged 22-36 (25.6±2.5)) participated in the present study. The Soroka Medical Center Internal Review Board approved the experimental procedures and written informed consent was obtained from each subject. The sample size was selected so that the correlation effect size of 0.4 would have power greater than 1 − β = 0.75 (one-tailed test), with α set to 0.05. According to G*Power (Faul et al., 2009), the required minimum sample size is 30.

### Experimental Setup and Design

Subjects lay in the scanner bore and viewed a back-projected screen through an angled mirror, which prevented any visual feedback of their arm and hand. An MRI-compatible digitizing tablet (Hybridmojo LLC, CA, USA) was placed over the subject’s waist and used to track their arm movements (Figure 1A). Subjects performed slice (out-and-back) reaching movements from a central target to four peripheral targets differing in their directions and extents (Figure 1B) and did not receive any visual feedback of their arm location during movement. Each trial started with the presentation of a peripheral target for one second. Four seconds after the target disappeared, the central target changed from red to green, indicating that the movement should be performed by moving the stylus pen on the tablet. Subjects had one second to complete the movement after which the center target turned red and remained red for the entire intertrial-interval (ITI), which lasted six seconds. There was no post-trial visual feedback or knowledge-of-results. All subjects preformed three experiments with each arm, each lasted 9 minutes and contained 11 movements to each of the four targets in a random order.

### Movement Recording and Analysis

Kinematic data were recorded at 200 Hz. Trials with a reaction time of more than 1 second, trials with a movement angle error >30° (at peak velocity or end point), and trials with movement length that was <50% or >200% of the target distance were discarded from further analysis. Trials containing correction movements (i.e., velocity profiles with more than two peaks) were also removed. On average approximately 8% (std 3%) of the trials were discarded for each subject. There was no significant difference in the number of discarded trials between the two arms.

We quantified intertrial variability for each of three kinematic components: movement direction, movement extent, and peak movement velocity. Movement extent and peak velocity variabilities were normalized by their respective means so as to compute the coefficient of variation (CV). This was necessary, because the variability of these kinematic components scales with their mean (speed-accuracy trade-off; Schmidt et al., 1979). Movement direction variability was quantified by the standard deviation (SD) across trials. Each of these measures was computed for each target and each subject separately and then averaged across targets to compute a single extent, peak velocity, and direction variability measure for each subject.

### MRI acquisition and preprocessing

Imaging was performed using a Philips Ingenia 3T MRI scanner located at the Ben-Gurion University Brain Imaging Research Center. The scanner was equipped with a 32 channel head coil, which was used for RF transmit and receive. Blood oxygenation level-dependent (BOLD) contrast was obtained using a T2* sensitive echo planar imaging (EPI) pulse sequence (TR = 2000 ms; TE = 35 ms; FA = 90^o^; 28 slices; voxel size of 2.6*2.6*3 mm and with 0.6 mm gap). Anatomical volumes were acquired with a T1-weighted sagittal sequence (TR = 8.165 ms; TE = 3.74 ms; FA = 8^o^; voxel size of 1*1*1 mm).

MRI data were preprocessed with the Freesurfer software package (http://surfer.nmr.mgh.harvard.edu, Fischl, 2012) and FsFast (Freesurfer Functional Analysis Stream). Briefly, this process includes removal of non-brain tissue and segmentation of subcortical, gray, and white matters based on image intensity. Individual brains were registered to a spherical atlas which utilized individual cortical folding patterns to match brain geometry across subjects. Each brain was then parcellated into 148 cortical ROIs using the Destrieux anatomical atlas (Destrieux et al., 2010). Functional scans were subjected to motion correction, slice-timing correction and temporal high-pass filtering with a cutoff frequency of two cycles per scan. Functional scans were registered to the high-resolution anatomical volume. No additional spatial smoothing was performed. Preprocessed data was imported into MATLAB (R2015a, *MathWorks Inc.* USA), and all further analysis was performed using custom software written in matlab.

### Time course analysis

To ensure that our estimates of intertrial fMRI variability were not generated by head motion, respiration, and blood flow artifacts, we removed the following components from the fMRI time-course of each cortical voxel, through linear regression: (1) six head motion parameters obtained by rigid body correction of head motion (three translations and three rotations), (2) fMRI time-course from the lateral ventricles, and (3) the mean fMRI signal of the entire cortex (i.e., global component). In addition, we normalized the time-course of each voxel to a mean of zero and unit variance (i.e., Z-score). This ensured that overall time-course variance was equal across subjects such that our measure of inter-trial fMRI variability captured only task-related trial-by-trial variability differences across subjects rather than variability associated with the entire scanning session.

### Identification of regions of interest

Visual and motor regions of interest (ROIs) were defined a priori according to a combination of anatomical and functional criteria in the native space of each subject. We identified commonly reported visual, visuomotor, and motor ROIs by selecting 100 continuous functional voxels with the strongest activation (all movements > rest) in each of the following anatomical locations: Early visual cortex (Vis) - Occipital pole and calcarine sulcus; Superior parietal lobule (SPL) - Anterior portion of the superior parietal lobule, superior to the IPS and slightly posterior to the postcentral sulcus; Inferior parietal lobule (IPL) - Dorsal portion of the angular gyrus and the middle segment of the intraparietal sulcus; Primary motor cortex (M1) - anterior bank of the central sulcus in the hand knob area; Dorsal premotor cortex (PMd) - Junction of superior frontal sulcus and precentral sulcus; Ventral premotor cortex (PMv) - Junction of inferior frontal sulcus and precentral sulcus; and Supplementary motor area (SMA) - Medial wall of the superior frontal gyrus, anterior to the central sulcus, posterior to the vertical projection of the anterior commissure.

We also defined control ROIs that did not exhibit task-related activations in the dorsolateral prefrontal cortex (dlPFC) - middle frontal sulcus, and 8 ROIs located outside the brain/head of the subject (one ROI in each corner of the scanned volume). These control ROIs enabled us to demonstrate the specificity of the results to the visuomotor cortical ROIs.

### Intertrial fMRI variability

Variability across trials was computed for each subject separately, relative to their mean hemodynamic response in each ROI. We computed a hemodynamic response function (HRF) for each target and then built a general linear model (GLM) with a row for every time-point and a column for every trial. Each column contained a delta function at trial onset, which was convolved with the relevant target-specific HRF. This enabled us to estimate a response amplitude (beta value) for each trial using multiple regression. Intertrial fMRI variability was estimated as the standard deviation across beta values (trials) to each of the targets. Before examining the correlations of individual fMRI variability magnitudes across targets and arms, we first regressed-out the subjects’ framewise displacement magnitudes. This ensured that individual fMRI variability measures were not generated by potential differences in head motion (Power et al., 2012).

Previous studies have reported that when using a canonical HRF to estimate intertrial variability, the strength of the mean fMRI response is positively correlated with intertrial variability (Ferri et al., 2015; He, 2013). This is problematic, because it does not allow for a clear separation of the two measures and their behavioral relevance. By quantifying intertrial variability with respect to each subject’s HRF, rather than using a canonical HRF, we were able to entirely discount the mean HRF amplitude and shape from our analysis – yielding a pure (isolated) measure of individual intertrial variability.

### Correlations

We used Pearson correlation coefficients to assess whether individual kinematic variability magnitudes were correlated across targets, arms, and different kinematic components. Equivalent analyses were performed to examine whether individual fMRI variability magnitudes (in each of the examined ROIs) were correlated across targets and arms as well as between the variability of each kinematic component and fMRI variability in each ROI. We assessed the statistical significance using a permutation tests. We randomly shuffled the variability values of the different subjects in each correlation analysis and computed the correlation. This process was repeated 5000 times to generate 5000 correlation values that represented a distribution of correlations expected by chance (null distribution). For the true (un-shuffled) value to be considered significant, it had to surpass the 97.5th percentile of the null distribution (i.e., the equivalent of a p < 0.05 value in a two-tailed t-test). We used the false discovery rate (FDR) correction (Benjamini and Hochberg, 1995; Yekutieli and Benjamini, 1999) to correct for the multiple comparisons across target pairs and across ROIs.

### Searchlight analysis

In addition to the ROI analysis, we used a searchlight analysis (Kriegeskorte et al., 2006) to map the correlations between fMRI variability and kinematic variability (i.e., movement extent, peak velocity, or direction) throughout the entire cortex. Clusters of 125 functional voxels were defined using a cube with an edge length of 5 voxels around each gray matter voxel in the native space of each subject. fMRI variability was calculated for each cluster of voxels, as described above in the ROI analysis. After computing the variability map of each subjects, all maps were transformed to a standard cortical surface using Freesurfer, and correlation analysis between kinematic and fMRI variabilities were performed for each kinematic measure using movements performed by either right or left arm. This yielded six correlation maps (three kinematic variables and two arms). A student t-test was used to determine the significance of the correlation across subjects in each vertex. We used FDR correction to correct for the multiple comparisons performed across vertices (Storey, 2002).

## Acknowledgements

We would like to thank Ilan Shelef and Moti Salti for their help in acquiring the fMRI data, and Lior Shmuelof for helpful discussions about the manuscript. The research described in this paper was supported by ISF grant 961/14 (I.D.), Helmsley Foundation (O.D.) and the ABC Robotics Center.

